# Development of optogenetic tools to manipulate cell cycle checkpoints

**DOI:** 10.1101/2020.06.22.166264

**Authors:** Yuhei Goto, Kazuhiro Aoki

## Abstract

In order to understand the systematic regulation of the cell cycle, we need more precise tools for cell-cycle perturbation. Optogenetics is a powerful technique for precisely controlling cellular signaling at higher spatial and temporal resolution. Here, we report optogenetic tools for the rapid and reversible control of cell-cycle checkpoints with a red/far-red light photoreceptor, phytochrome B (PhyB). We established fission yeast cells producing phycocyanobilin as a chromophore of PhyB, and demonstrated light-dependent protein recruitment to the plasma membrane, nucleus, and kinetochore. Using this system, we developed optogenetic manipulation of the cell cycle in two ways: the Opto-G2/M checkpoint triggered G2/M cell cycle arrest in response to red light, and Opto-SAC induced a spindle assembly checkpoint (SAC) in response to red light and then quickly released the SAC by far-red light.

## INTRODUCTION

The cell cycle is a highly robust and regulated biological process that ensures the propagation of genetic information to daughter cells. Chromosome DNAs are replicated during the S phase and precisely divided into daughters in the M phase. A huge number of biochemical events accompany the cell cycle progression, and these are fundamentally regulated through CDK-cyclin, a master regulator kinase complex of the cell cycle. It is widely believed that different CDK-cyclin complexes are sequentially formed in each cell cycle phase, thereby phosphorylating distinct substrates in the distinct cell cycle phase specificities (Bhaduri and Pryciak, 2011; Kõivomägi et al., 2011; Loog and Morgan, 2005; Pagliuca et al., 2011). Meanwhile, it has been reported that CDK-cyclin complexes are exchangeable and that substrate-specific thresholds of CDK activity suffice to determine the cell cycle phase-specific phosphorylation (Swaffer et al., 2016). In fission yeast, only a single CDK-cyclin complex is sufficient for the progression of the cell cycle (Coudreuse and Nurse, 2010), and thus fission yeast can be used to investigate the general mechanisms of the CDK-cyclin complex in cell cycle progression. For example, fission yeast has been used to artificial manipulate CDK-cyclin activity; the temperature-sensitive mutants of the CDK-cyclin regulatory network were originally identified as cell division cycle (*cdc*) mutants. Among them, *cdc25-ts* has since been well established as a mutant capable of reversibly controlling the cell cycle (Moreno et al., 1989). In addition, chemogenetic manipulation of CDK activity such as ATP analog sensitive mutants also enables rapid and reversible inhibition of CDK-cyclin activity, thereby controlling cell cycle progression (Aoi et al., 2014; Dischinger et al., 2008). However, these methods are still limited by technical difficulties in rapidly and repeatedly changing the temperature or medium, and only a few alternative methods for manipulating CDK-cyclin activity have been developed (Niopek et al., 2014).

The spindle assembly checkpoint (SAC) (also known as the mitotic checkpoint) is an evolutionarily conserved safeguard mechanism by which chromosomes faithfully segregate at anaphase (London and Biggins, 2014; Musacchio and Salmon, 2007). The SAC delays anaphase onset until all chromosomes are properly attached to the spindle microtubule. The core SAC machinery is composed of Mad1, Mad2, Bub1, Bub3, BubR1/Mad3, and Mps1/Mph1 in fission yeast (Yamagishi et al., 2014). Mps1/Mph1 is the most upstream component of SAC; Mps1/Mph1 binds to the outer kinetochore through the Ndc80 complex, and competes for the binding of microtubule to the Ndc80 complex (Hiruma et al., 2015; Ji et al., 2015). The Mps1/Mph1 that is enriched at unattached kinetochores emits a “wait anaphase” signal through the accumulation of downstream SAC proteins. This hierarchical regulation by Mps1/Mph1 enables artificial activation of SAC by tethering Mps1/Mph1 at the kinetochore through fusion with the kinetochore scaffold (Heinrich et al., 2012; Ito et al., 2012; Jelluma et al., 2010; Yamagishi et al., 2012). In mammalian cells, forced localization of Mad1 also activates SAC ectopically (Maldonado and Kapoor, 2011). These artificial activations of SAC demonstrate that the localization of Mps1/Mph1 at the kinetochore is sufficient for SAC activation. Several research groups have reported chemical-induced dimerization (CID) systems for artificial SAC activation in fission yeast and mammalian cells (Amin et al., 2018; Ballister et al., 2014; Chen et al., 2019). However, the chemical treatments are limited to reversibility and spatio-temporal regulation. Despite the decades of SAC studies, it still remains unclear how long cells can arrest their cell cycles through SAC with competence for the next cell cycle.

Optogenetics is a promising tool to control intracellular signaling in a rapid and reversible manner with high spatial resolution. Lampson and his colleagues developed optogenetic tools for controlling kinetochore functions including artificial SAC activation and silencing with a combination of chemical biology and optogenetics (Aonbangkhen et al., 2018; Zhang et al., 2017). Thus, a fully genetically encoded optogenetics tool controlling SAC has not yet been developed. Here, we report fully genetically encoded systems for optogenetic control of the cell cycle in fission yeast, which we call Opto-G2/M checkpoint and Opto-SAC. These systems rely on the rapid and reversible heterodimerization of proteins in response to light. We exploit these light-inducible dimerization (LID) systems for protein translocation to modulate CDK activity and SAC switching, revealing that SAC and cytokinesis checkpoint are independently regulated.

## RESULTS

### Development of a fully genetically encoded PhyB-PIF LID system in fission yeast

To achieve rapid and reversible manipulation of the cell cycle, we focused on a phytochrome B (PhyB)-phytochrome interacting factor (PIF) LID system. The heterodimerization between PhyB and PIF is quickly triggered on and off by exposure to 635 nm and 730 nm light (Levskaya et al., 2009) (Fig. 1A). The red/far-red photoreceptor, PhyB, requires a chromophore, i.e., phytochromobilin and phycocyanobilin (PCB), for the photoreception (Kami et al., 2004; Kohchi et al., 2001). However, these linear tetrapyrroles do not exist in non-photosynthetic organisms such as yeast and mammalian cells, which makes it difficult to use the PhyB-PIF LID system. We and other groups have overcome this issue by biosynthesis of PCB in mammalian cells and fission yeast (Kyriakakis et al., 2018; Müller et al., 2013; Uda et al., 2017); PCB biosynthesis is induced by the expression of four proteins (HO1, PcyA, Fd, and Fnr) in the mitochondria, where heme is abundant (Fig. 1B and 1C). In our PCB synthesis system, hereinafter referred to as SynPCB, we exploit a polycistronic synthetic gene in which these four genes are linked by a P2A peptide sequence. Recently, we have succeeded in improving this SynPCB system (Uda et al., 2020), but, in this study, the original SynPCB1.0 was used for fission yeast.

**Figure 1.**
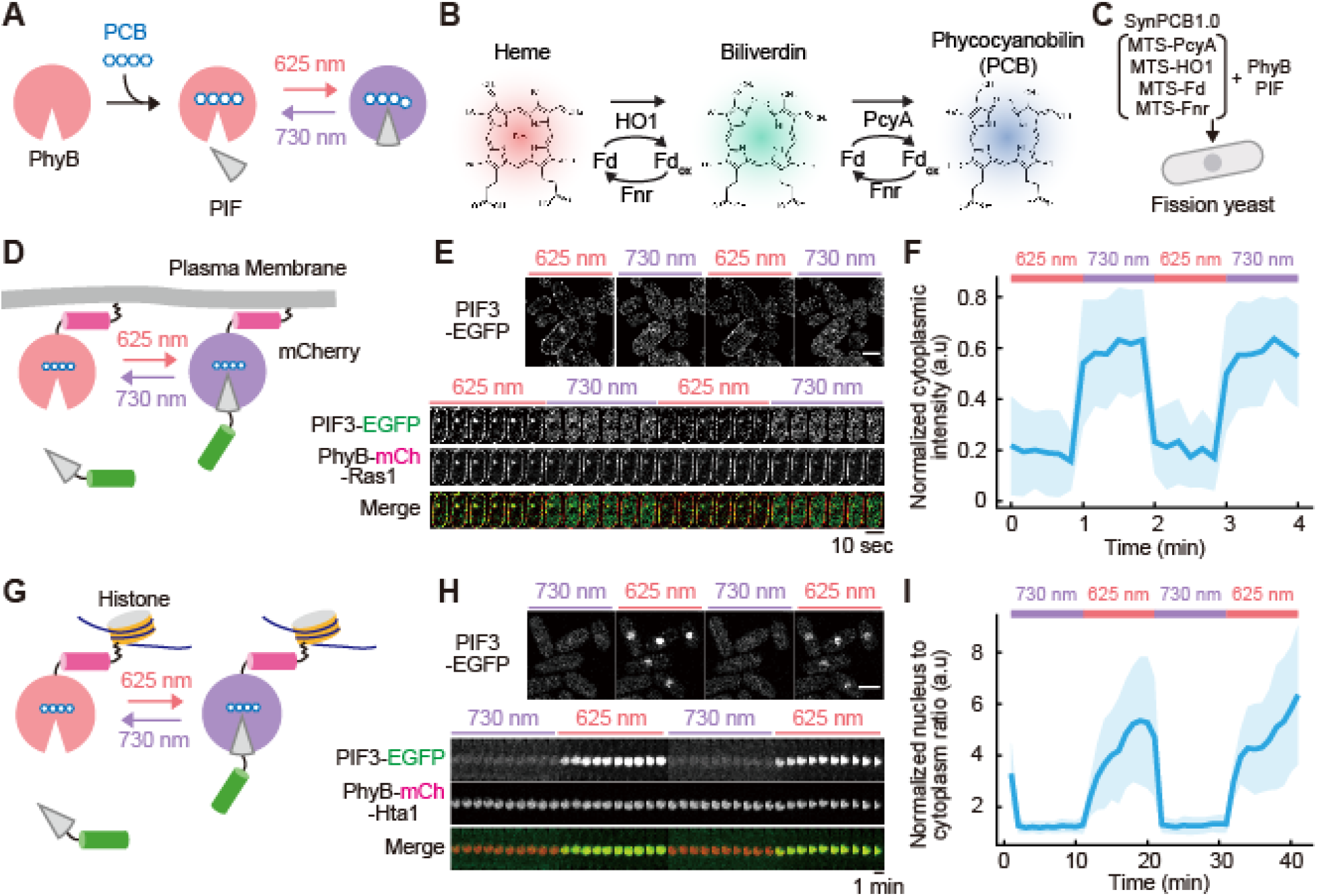
Genetically encoded PCB biosynthesis and PIF3 recruitment to PhyB in *Schizosaccharomyces pombe*. (A) Light-inducible dimerization (LID) of PhyB and PIF. (B) Phycocyanobilin (PCB) synthesis pathway in cyanobacteria. (C) Schematic representation of the genetically encoded PhyB-PIF LID system, in which PCB biosynthesis is induced by SynPCB1.0 comprised of mitochondria targeting sequence (MTS)-fused PcyA, HO1, Fd, and Fnr. (D) Schematic of light-induced recruitment of PIF3-EGFP to PhyB-mCh-Ras1 at the plasma membrane. (E) Representative fluorescence images of PIF3-EGFP recruitment to PhyB-mCh-Ras1 (upper). Kymograph of repeated PIF3-EGFP recruitment to PhyB-mCh-Ras1 at the plasma membrane with a time interval of 10 sec (lower). Images were obtained by confocal laser scanning microscopy. Scale bars, 5 μm. (F) Membrane-recruited PIF-EGFP was quantified by the change in the fluorescence intensity of PIF-EGFP at the cytoplasm. The average values (bold lines) are plotted as a function of time with the SD. N = 10. (G) Schematic of light-inducible recruitment of PIF3-EGFP to PhyB-Hta1 at the nucleus. (H) Representative fluorescence images of PIF3-EGFP recruitment to PhyB-mCh-Hta1 (upper). Kymograph of repeated PIF3-EGFP recruitment to PhyB-mCh-Ras1 at the nucleus with a time interval of 1 min (lower). Images were obtained by confocal laser scanning microscopy. Scale bars, 5 μm. (I) Nuclear-recruited PIF-EGFP was quantified by the change in the fluorescence intensity of PIF-EGFP at the nucleus. The average values (bold lines) are plotted as a function of time with the SD. N = 15.

First, we compared PCB synthesis by the SynPCB system and external PCB uptake in fission yeast. PCB was quantified by using the PhyB-Y276H mutant, which emits near-infrared fluorescence when bound to PCB (Fig. S1A). Consistent with the previous study, the expression of SynPCB1.0 increased near-infrared fluorescence from the PhyB-Y276H mutant (Fig. S1B). In contrast, the uptake of external purified PCB required a high dose and a treatment time longer than the duration of the fission yeast cell cycle (~ 2 h) (Fig. S1C). We did not observe any growth defect of fission yeast by overexpression of SynPCB1.0 under normal growth conditions (Fig. S1D and S1E).

Next, we confirmed the light-induced association and dissociation of PhyB-PIF in fission yeast by visualizing the changes in subcellular localization of the protein. The PhyB (a.a. 1-621) tagged with mCherry and CAAX sequence of fission yeast Ras1 (a.a. 201-260) at the C-terminus (PhyB-mCh-Ras1) was localized at the plasma membrane, thereby acting as a localizer (Fig. 1D and 1E). PIF3 (a.a. 1-100) fused with EGFP (PIF3-EGFP) was repeatedly and quickly translocated between the plasma membrane and cytosol upon red light (625 nm) and far-red light (730 nm) exposure through the association and dissociation of PhyB-mCh-Ras1 (Fig. 1E and 1F, Movie S1). As another example, the light-induced nucleocytoplasmic shuttling of PIF3-EGFP was implemented by using the histone-targeted PhyB-mCherry (*PhyB-mCh-hta1*) (Fig. 1G). As expected, PIF3-EGFP accumulated at the nucleus in response to red light exposure, and this accumulation was released by far-red light exposure (Fig. 1H and 1I). It should be noted that the nuclear translocation of PIF3-EGFP showed slower kinetics than plasma membrane translocation (Fig. 1H and I), possibly due to the slow nuclear import of PIF3-EGFP. In contrast, in both cases, far-red light caused very rapid dissociation of PIF3-EGFP from PhyB. These data lead to the conclusion that our genetically encoded PhyB-PIF LID system can be used for the rapid and reversible translocation of proteins in fission yeast.

### Opto-G2/M checkpoint: Optogenetic manipulation of the G2/M checkpoint

To manipulate the cell cycle progression by optogenetics, we focus on the cyclin-dependent kinase (CDK) in fission yeast. Fission yeast has a single CDK, Cdc2, for the mitotic cell cycle, and Cdc2 activation is known to be required for the G2/M cell-cycle checkpoint progression (Dischinger et al., 2008). Cdc2 is mainly localized at the nucleus through its association with cyclin B, Cdc13p (Decottignies et al., 2001) To achieve this model, endogenous Cdc2 was tagged with PIF3 and EGFP (*cdc2-PIF3-EGFP*) to exclude Cdc2 from the nucleus through the binding to PhyB-mCh-Ras1 at the plasma membrane by red light (Fig. 2A). We named this system Opto-G2/M checkpoint. The cells expressing Cdc2-PIF3-EGFP with PhyB-mCh-Ras1 and SynPCB1.0 stopped proliferation upon the illumination of red light, whereas the cells illuminated by far-red light grew well (Fig. 2B). Similarly, endogenous Cdc25, the CDK activating phosphatase, was also tagged with PIF3 and EGFP (*cdc25-PIF3-EGFP*) to exclude Cdc25 from the nucleus, and these cells demonstrated a growth defect in a light-dependent manner (Fig. 2B). Upon far-red light illumination, the cells grew properly with normal morphology and nuclear accumulation of Cdc2-PIF3-EGFP (Fig. 2C). After the incubation of this strain with red light illumination, many cells exhibited elongated cell bodies and nuclei, which are the typical phenotypes of CDK inhibition in fission yeast (Fig. 2C). Part of Cdc2-PIF3-EGFP was trapped on the plasma membrane, but the majority was still retained in the nucleus, suggesting that partial inhibition of CDK activity is sufficient to block the cell cycle progression in fission yeast.

**Figure 2.**
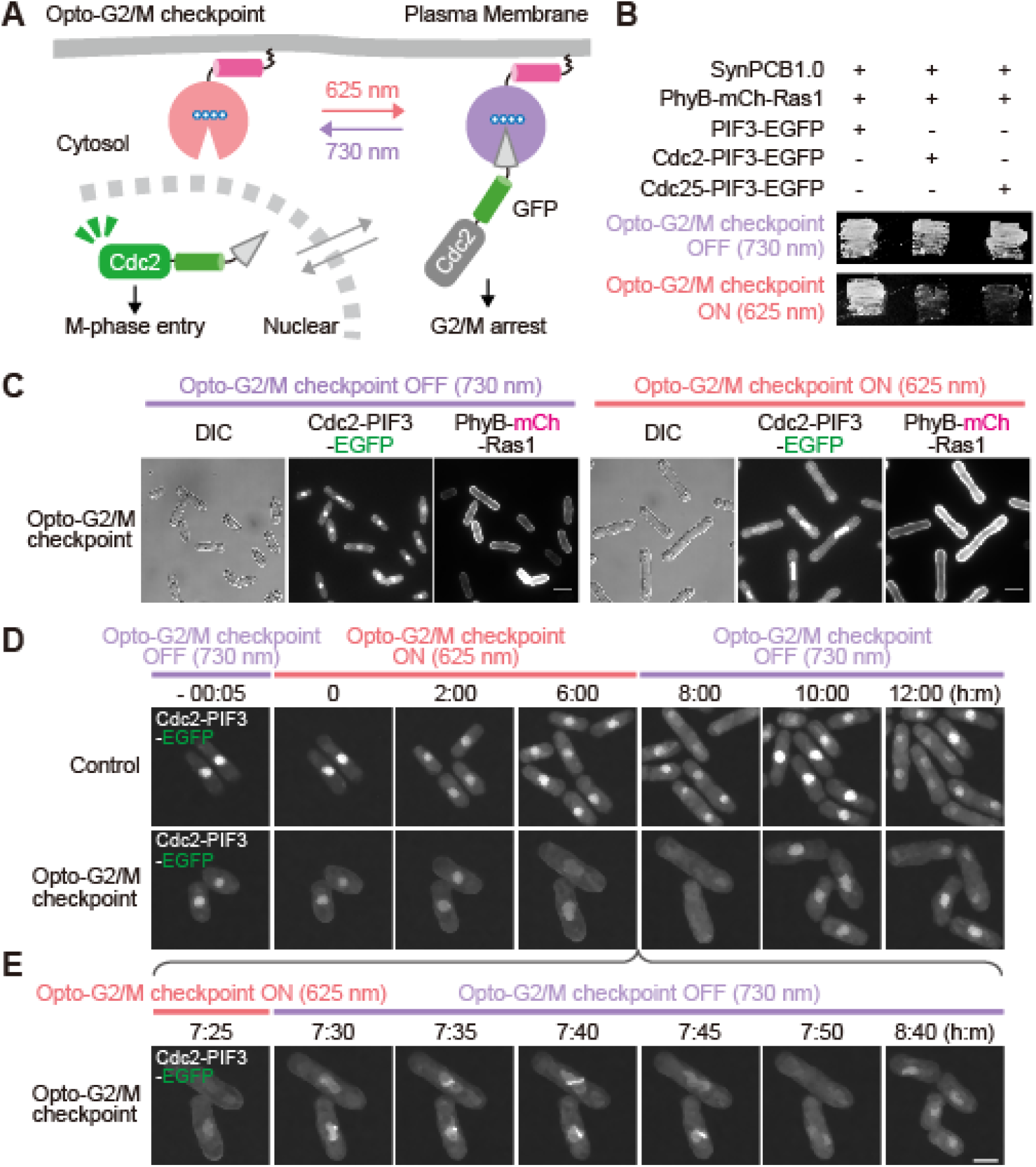
Optical manipulation of cell cycle progression through light-inducible control of CDK activity (the Opto-G2/M checkpoint). (A) Schematic of the design of the Opto-G2/M checkpoint. Upon red light illumination, Cdc2-PIF3-EGFP is exported from the nucleus through the binding to membrane-targeted PhyB-mCh-Ras1. (B) Fission yeast expressing the indicated components were exposed to either far-red light (upper) or red light (lower), showing that the proliferation of cells expressing the Opto-G2/M checkpoint (Cdc2 or Cdc25) was inhibited in a light-dependent manner. (C) Representative images of Opto-G2/M checkpoint-expressing fission yeast cells exposed by far-red nm light (upper) and red nm light (lower), respectively. Fluorescence images were obtained by wide-field fluorescence microscopy. Scale bars, 5 μm. (D and E) Cells expressing all components of the Opto-G2/M checkpoint (lower) and control cells lacking PhyB-mCh-Ras1 (upper) were imaged by spinning disk confocal microscopy. Representative images of endogenous Cdc2-PIF3-EGFP are shown. The Opto-G2/M checkpoint was switched ON and OFF by irradiating red and far-red light, respectively. In control cells, PhyB-mCh-Ras1 expression was suppressed by the addition of thiamine. Detailed images of the Opto-G2/M checkpoint upon switching from ON to OFF are shown (E). Scale bar, 5 μm.

We then assessed whether the Opto-G2/M checkpoint could reversibly control the G2/M checkpoint. The cells were arrested at the G2 phase by red light exposure, followed by far-red light irradiation to resume the cell cycle. Control cells lacking PhyB-mCh-Ras1 did not show any cell cycle arrest by light irradiation (Fig. 2D). The red light-induced G2 arrest and far-red light-induced resumption of the cell cycle were observed in the cells expressing all components required for the Opto-G2/M checkpoint (Fig. 2D and Movie S1). Cdc2-PIF3-EGFP trapped in the plasma membrane was rapidly translocated to the nucleus by far-red light irradiation, which in turn led to cell-cycle progression from G2 into mitosis (Fig. 2E). We should note that the cell cycle arrest by Opto-G2/M checkpoint is heterogeneous from cell to cell; that is, a mixture of cells is observed—cells in which the cell cycle stops and cells in which the cell cycle continues. This is probably due to the heterogenous sequestration of Cdc2 proteins.

### Opto-SAC: Optical control of the spindle assembly checkpoint

Next, we attempted to manipulate the spindle assembly checkpoint (SAC). SAC requires the hierarchical accumulation of checkpoint proteins at unattached kinetochores, and Mph1, which is the top of this hierarchy, causes ectopic SAC activation and cell cycle arrest at metaphase when it is artificially tethered at kinetochores (Heinrich et al., 2012; Ito et al., 2012; Yamagishi et al., 2012)(Fig. 3A). Therefore, we aimed to develop an optogenetically controllable SAC system (Opto-SAC), which achieves red light-inducible recruitment of Mph1 to kinetochores and far-red light-induced detachment of Mph1 from kinetochores (Fig. 3B). The advantage of this Opto-SAC system over other inducible SAC systems is that it can rapidly induce both recruitment and detachment of Mph1 by light.

**Figure 3.**
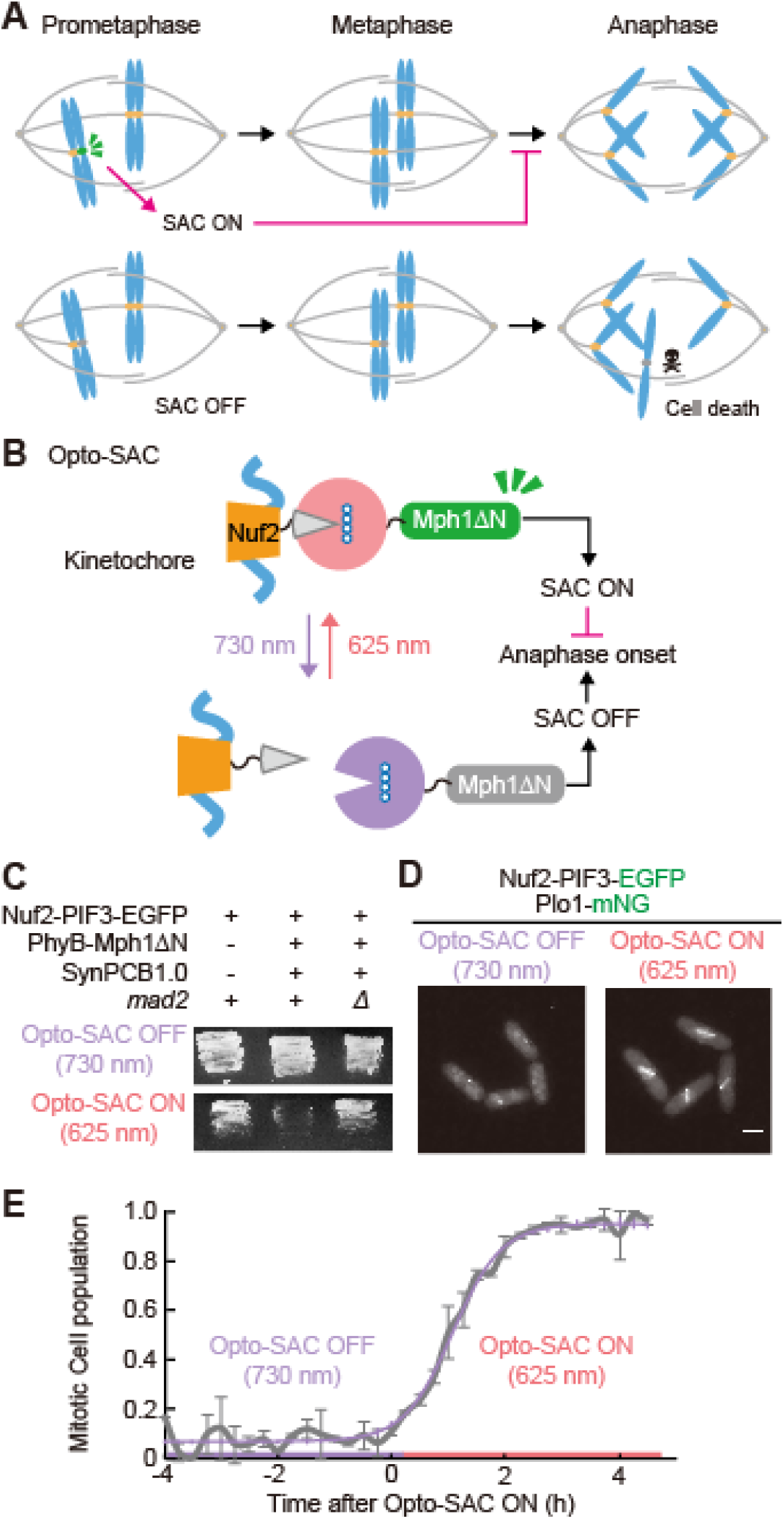
Light-inducible control of the spindle assembly checkpoint (Opto-SAC). (A) Schematic of the spindle assembly checkpoint (SAC), which ensures the faithful chromosome segregation and anaphase onset by monitoring kinetochore-microtubule attachment. (B) Schematic of the design of Opto-SAC. Mph1ΔN, which lacks the kinetochore binding domain, is fused with PhyB (*PhyB-mph1ΔN*). Endogenous Nuf2 is fused with PIF3 (*nuf2-PIF3*) as a kinetochore localizer. Upon red light illumination, PhyB-Mph1ΔN is recruited to kinetochores through the binding to Nuf2-PIF3, and SAC turns ON. Far-red light exposure induces dissociation of PhyB-Mph1ΔN from kinetochores, leading to the inhibition of activated SAC. (C) Fission yeast expressing the indicated components was exposed to either far-red light (upper) or red light (lower), showing that the proliferation of cells expressing Opto-SAC was inhibited in a SAC-dependent manner. (D) Representative fluorescence images of cells under Opto-SAC OFF (left) and Opto-SAC ON (right) conditions. Images were obtained by wide-field fluorescence microscopy. Scale bars, 5 μm. (E) The fraction of mitotic cells is plotted as a function of time after turning Opto-SAC ON (N > 80 cells at each time point).

To implement Opto-SAC, endogenous Nuf2, a kinetochore complex protein, or Mis12, an outer kinetochore protein, was tagged with PIF3 as a kinetochore localizer (Fig. 3B and Fig. S2). As an actuator, PhyB was fused with the C-terminal kinase domain of Mph1 (*PhyB621-mph1ΔN*) lacking an authentic kinetochore localization domain (Heinrich et al., 2012)(Fig. 3B). PhyB-Mph1ΔN was exogenously expressed under the weak constitutive promoter. We examined cell proliferation under red or far-red light illumination conditions. As we expected, continuous red light exposure, i.e., the Opto-SAC ON state, inhibited cell proliferation (Fig. 3C). Importantly, the deletion of *mad2*, a downstream effector of the SAC signaling cascade, rescued the cell proliferation even under an Opto-SAC ON condition (Fig. 3C), indicating that Opto-SAC indeed triggers SAC in a light-dependent manner.

Next, we tried to confirm that cells were indeed arrested at metaphase in the Opto-SAC ON state. To this end, the endogenous *plo1* gene was tagged with mNeonGreen (mNG) as the marker for mitotic cells. In the Opto-SAC OFF state under which far-red light was irradiated, only a few cells were in the mitosis phase (Fig. 3D, left). After turning on the Opto-SAC by red light exposure, most cells demonstrated aligned kinetochores decorated with Nuf2-PIF3-EGFP and Plo1-mNG, clearly indicating mitotic arrest (Fig. 3D, right). We quantified the dynamics of the fraction of mitotic cells after turning on the Opto-SAC. Under the Opto-SAC OFF condition, less than 10% of the cells were mitotic on average (Fig. 3E). The fraction of mitotic cells increased after turning on the Opto-SAC, and almost all cells underwent mitotic arrest after 2 h (Fig. 3E). This time scale is consistent with the duration of the fission yeast cell cycle, implying that once cells enter mitosis under the Opto-SAC ON condition, they are arrested at metaphase without any defect in any other cell cycle stages.

### Continuous metaphase arrest by Opto-SAC reduces the competency for SAC silencing

Taking advantage of the Opto-SAC system, which is capable of rapidly inducing both SAC ON and OFF by light, we examined how cells arrested to metaphase are released from SAC and returned to the normal cell cycle. To visualize the mitotic exit, the endogenous cyclin B gene, *cdc13*, was tagged with mNeonGreen, which is accumulated in the nucleus as the cell cycle progress and degraded rapidly at the end of mitosis. When the SAC was turned on by red light, the cell cycle was arrested at metaphase, coincident with the nuclear accumulation of Cdc13-mNG (Fig. 4A). Subsequently, far-red light illumination induced the SAC OFF state, leading to the rapid degradation of Cdc13-mNG and mitotic exit in the majority of cells, but some cells either slowly returned to the normal cell cycle or remained arrested in the metaphase (Fig. 4A). Next, we investigated whether SAC was released by far-red light after the cells were subjected to red light irradiation of different durations to induce SAC. Interestingly, the longer the SAC ON state, the greater the fraction of cells with delayed reversion to the cell cycle, and these cells remained arrested in the metaphase (Fig. 4B and 4C). We found that the cells in which SAC silencing failed showed a continuous increase in the fluorescence intensity of Cdc13-mNG even when Opto-SAC was turned off by far-red light. The failure of SAC silencing did not correlate with the timing of cell division before turning on the Opto-SAC (Fig. S3). The induction of long-term SAC by Opto-SAC caused the spindle fragmentation and premature septation, which were typical Cut phenotypes, as visualized by Plo1-mNG and Nuf2-PIF3-EGFP (Fig. 4D and E), suggesting an uncoupling of the SAC and cytokinesis checkpoints.

**Figure 4.**
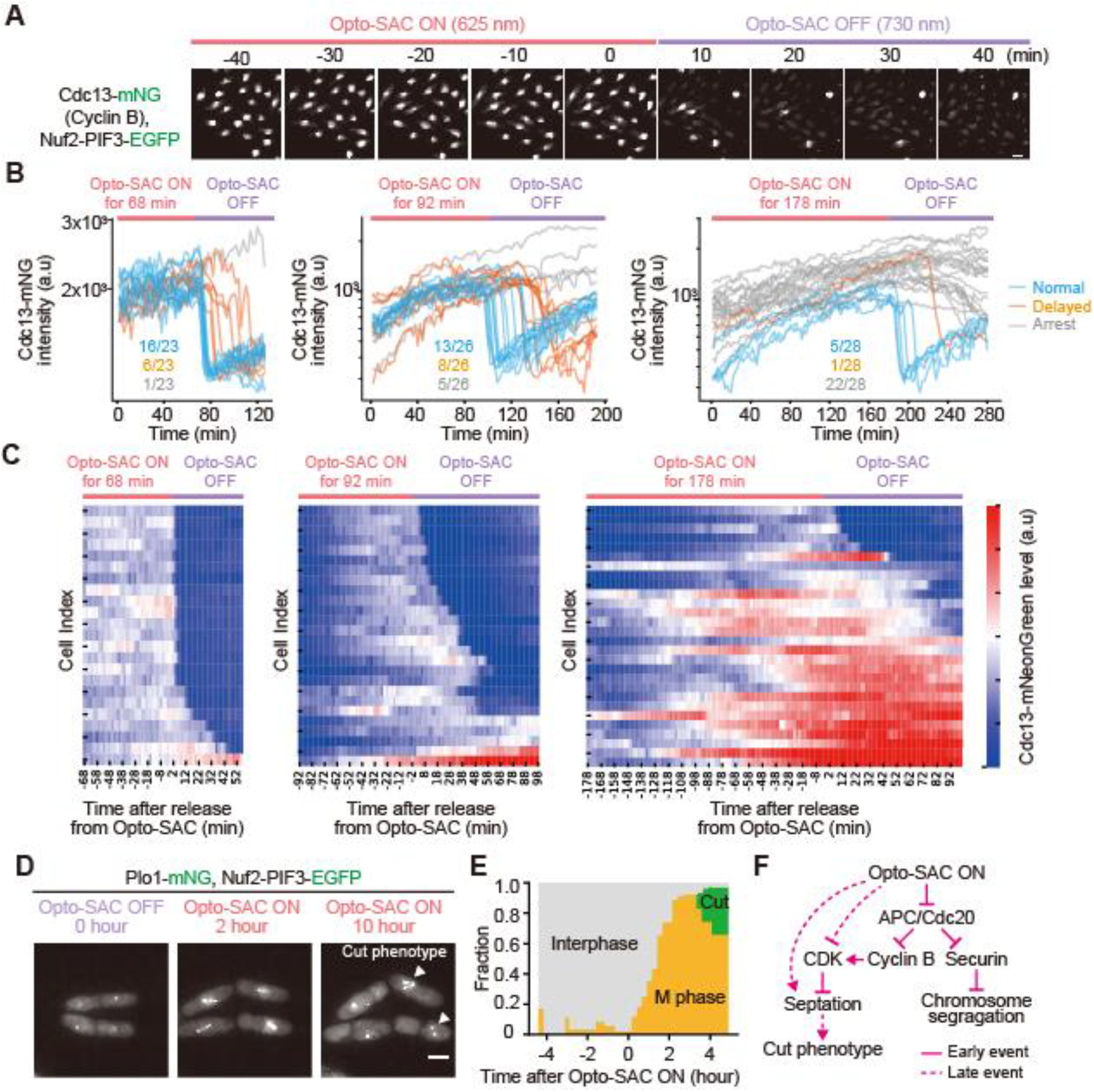
Long-term SAC activation leads to premature septation arrest without cyclin B degradation. (A) The cells were arrested at metaphase by turning Opto-SAC ON, followed by induction of Opto-SAC OFF by far-red light exposure. Representative images of endogenous cyclin B (Cdc13-mNG) and Nuf2-PIF3-EGFP are shown. Images were obtained by wide-field fluorescence microscopy. Scale bars, 5 μm. (B) The fluorescence intensity of cyclin B (Cdc13-mNG) is plotted as a function of time after Opto-SAC OFF. Blue and orange lines indicate the cells showing the rapid and delayed degradation of Cdc13-mNG after the Opto-SAC OFF, respectively. Gray lines are the cells that never underwent Cdc13-mNG degradation (N = 23, 26, and 28, respectively). (C) Heatmaps of the time course of cyclin B (Cdc13-mNG) fluorescence intensity around the Opto-SAC OFF are shown. (D) Representative cells undergoing a failure of mitosis under long-term metaphase arrest by Opto-SAC. Images were obtained by wide-field fluorescence microscopy. Scale bars, 5 μm. (E) Time course of the fraction of cells in interphase, M phase, and cut phenotype before and after Opto-SAC ON. (F) Schematic of the working model.

## DISCUSSION

In this study, we developed two optogenetic tools, namely Opto-G2/M checkpoint and Opto-SAC, for cell cycle control in a fission yeast, *Schizosaccharomyces pombe;* the former system inhibits Cdc2/CDK1 activity by red light, while the latter system induces SAC by red light, which can then be released by far-red light. A major factor for the success of the PhyB-PIF LID system in fission yeast is the sufficient biosynthesis of PCB without any growth defect (Fig. S1). The level of PCB biosynthesis is probably adequate because the fission yeast genome does not encode a biliverdin reductase gene, which degrades PCB and generates phycocyanorubin (Terry et al., 1993; Uda et al., 2017).

Opto-SAC allows us to rapidly and reversibly control the activation of SAC by light. Several SAC activation tools using small chemical compounds and CID systems have been reported so far. Although these tools can easily induce SAC activation, it is difficult to rapidly restore the activated SAC. We found that M phase arrest by Opto-SAC for a long duration induced premature septation and spindle collapse, leading to the cut phenotype (Fig. 4). Surprisingly, by tracing the cell cycle progression at the single-cell level, we observed that some cells arrested by Opto-SAC undergo cytokinesis without any cyclin B Cdc13-mNG degradation. This phenotype is clearly distinct from the previously reported mitotic slippage, which evades mitotic arrest and proceeds to interphase through cyclin B degradation during the activation of SAC (Bonaiuti et al., 2018; Brito and Rieder, 2006). In these studies, the SAC activation was achieved by microtubule destabilizers such as nocodazole. On the other hand, Opto-SAC affects only the downstream Mph1 signaling pathway. These results suggest that long-term SAC leads to premature septation directly or indirectly through CDK inhibition (Fig. 4F), while microtubule destabilization induces unknown signaling that inhibits premature septation in addition to activating the SAC, but subsequently causes mitotic slippage with cyclin B degradation.

Opto-G2/M checkpoint suppresses cell cycle progression by forced localization of Cdc2 to the plasma membrane. This requires the expression of a sufficient amount of the localizer, PhyB-mCh-Ras1, and labeling of the endogenous *cdc2* gene with the *PIF3* gene. Although the subcellular localization of CDK-cyclin and its regulation differ among organisms, forced sequestration of CDKs from their authentic intracellular localization is a promising strategy for CDK inactivation in other systems. Indeed, similar approaches have been reported to control CDK activity by optogenetics (Niopek et al., 2014; Yang et al., 2013). It is difficult to activate CDK activity in our Opto-G2/M checkpoint system, although this limitation can be overcome by introducing opto-kinase approaches showing light-induced conformational change (Leopold et al., 2018; Zhou et al., 2017). Compared to the Opto-G2/M checkpoint system, the Opto-SAC system has several advantages: the Opto-SAC system can be technically implemented by exogenously expressing two components (PhyB-kinetochore localizer and PhyB-Mph1ΔN), making it easy to apply to other experimental systems such as mammalian cultured cells. Further, since SAC is highly sensitive to Mph1 activity, only a small number of Mph1ΔN proteins suffices to induce SAC. Thus, we believe that Opto-SAC does not require stringent control of the PhyB-PIF expression ratio than that should be achieved by the Opto-G2/M checkpoint.

In summary, we report for the first time light-inducible control of protein translocation and the cell cycle with the PhyB-PIF LID system in fission yeast. These techniques would be useful for other experiments to optically modulate protein localization and activity not only in fission yeast but also in other species. The principal mechanisms underlying the CDK-mediated cell cycle progression and SAC are highly conserved among eukaryotes, and therefore Opto-G2/M checkpoint and Opto-SAC could be applied to other eukaryotes with minimal modification.

## Materials and Methods

### Fission yeast *Schizosaccharomyces pombe* strain and culture

All strains made and used in this study are listed in Table S1. The growth medium, sporulation medium, and other techniques for fission yeast were based on the protocol described previously (Moreno et al., 1991) unless otherwise noted.

### Plasmids

The cDNA of *Arabidopsis thaliana* PhyB (1–621 aa) was synthesized with codon optimization for fission yeast by gBlocks Gene Fragments (Integrated DNA Technologies). The cDNAs of *Arabidopsis thaliana* PIF3 (1–100 aa) and SynPCB1.0 were derived from pCAGGS-PIF3-linker-EGFP (Uda et al., 2017, 2020) by KOD One (TOYOBO). The cDNA of PhyB was inserted into the pREP1 or pNATZA21 backbone with *ras1, hta1*, or *mph1ΔN* genes cloned from the WT fission yeast genome. Endogenous tagging and deletion of genes were done as previously described (Moreno et al., 1991; Uda et al., 2017). The license of mNeonGreen was purchased from Allele Biotechnology & Pharmaceuticals, and mNeonGreen gene was synthesized with codon optimization for fission yeast by gBlocks Gene Fragment.

### Measurement of growth rate

Fission yeast cells were pre-cultured at 30 °C up to OD (600 nm) 1.0, following dilution to 1:100. A Compact Rocking incubator biophotorecorder TVS062CA (Advantec, Japan) was used for culture growth (30 °C, 70 rpm) and OD 660 nm detection. Growth curves were fitted by the logistic function (*x*= *K* / (1 + (*K/x_0_ - 1)e^-rt^*)) and doubling time (*ln2/r*) was calculated on Python 3 and Scipy.

### Live-cell fluorescence imaging

For the imaging with microfluidics, mid-log phase fission yeast cells in YEA were placed into a Y04C-02 microfluidic chamber controlled by a CellASIC Onix2 system (CAX2-S0000; Millipore). 300 μl YEA was placed into the flow-wells, the chambers were loaded with cells at 9 psi, and media were continuously flowed at 1 psi while heating at 32 °C by a temperature controllable manifold (CAX2-MXT20; Millipore). For confocal imaging, the cells were concentrated by centrifugation at 3,000 rpm, mounted on a slide glass, and sealed by a cover glass (Matsunami).

Widefield epifluorescence imaging was performed with an inverted microscope IX83 (Olympus) equipped with an oil immersion objective lens (UPlanSApo 60×/1.35; Olympus), sCMOS camera (Prime, Photometrics), and Spectra-X light engine (Lumencor Inc.). The microscope was controlled by MetaMorph software (Molecular Devices). For confocal fluorescence imaging, cells were imaged with a TCS SP5 microscope (Leica Microsystems) equipped with an oil immersion objective lens (HCX PL APO 63/×1.4–0.6 oil; Leica Microsystems). The excitation laser and fluorescence filter settings were as follows: excitation laser, 488 nm (mEGFP) and 633 nm (PCB fluorescence); excitation dichroic mirror, TD 488/543/633 dichroic mirror; detector, HyD 520–590 nm (mEGFP) and HyD 670–720 nm (PCB fluorescence). For the spinning disk confocal fluorescence imaging, cells were imaged with an IX83 inverted microscope equipped with the same camera as above, an oil objective lens (UPLXAPO 60×/1.42; Olympus) and a spinning disk confocal unit (CSU-W1; Yokogawa Electric Corporation) illuminated with a laser merge module containing 488 nm and 561 nm lasers. Excitation dichroic mirror: DM405/488/561; emission filters: 500–550 nm and 580–654 nm. LEDs for illumination with red (625 nm, LED-41VIS625; OptoCode) and far-red (735 nm, LED-41IR735; OptoCode) light were controlled manually.

### Imaging analysis

All fluorescence imaging data were analyzed and quantified in Fiji (Image J). The background was subtracted by the rolling-ball method. For the quantification of signal intensity, appropriate ROIs were manually selected. For the quantification of the mitotic cell population, cells having strong Plo1-mNG signals and aligned Nuf2-PIF3-EGFP were counted as mitotic cells, and cells having single Nuf2-PIF3-EGFP cells were counted as interphase cells.

### PCB treatment

PCB was purchased from Santa Cruz Biotechnology, dissolved in DMSO (final concentration, 5 mM), and stored at −30 °C. PCB was diluted by YEA, and cells were cultured for the indicated time in YEA liquid media containing the indicated concentration of PCB.

## Acknowledgments

We thank all members of the Aoki Laboratory for their helpful discussions and assistance. K.A. was supported by a CREST, JST Grant (JPMJCR1654), JSPS KAKENHI Grants (nos.16KT0069, 16H01425 “Resonance Bio”, 18H04754 “Resonance Bio”, 18H02444, and 19H05798), and the ONO Medical Research Foundation. Y.G. was supported by a JSPS KAKENHI Grant (no.19K16050) and a Jigami Yoshifumi Memorial Research Grant. Confocal images were acquired at the Spectrography and Bioimaging Facility of the NIBB Core Research Facilities.

## Author contributions

Y.G. and K.A. designed the research. Y.G. performed all experiments and data analysis. Y.G. and K.A. wrote the manuscript.

## Supplemental Information

**Figure S1. Comparison between biosynthesis and uptake of PCB in *Schizosaccharomyces pombe*.** (A) Schematic of the measurement of PCB fluorescence. Infra-red PCB fluorescence is observed when PCB binds to the PhyB mutant PhyB-Y276H.

(B) PCB fluorescence intensities in each cell are plotted as a function of mCherry fluorescence of PhyB-Y276H-mCherry in each cell. Upper and lower panels indicate control and SynPCB1.0-expressing cells, respectively.

(C) PCB fluorescence intensities in each cell are plotted as a function of mCherry fluorescence of PhyB-Y276H-mCherry. Cells expressing PhyB-Y276H-mCherry were incubated with the indicated concentration of PCB for 1 h (left) and 6 h (right). At least 110 cells were quantified in each graph.

(D) Growth curve of cells expressing SynPCB1.0. Solid lines indicate the mean OD from 3 independent cultures and shaded regions indicate the SD values.

(E) Doubling time was calculated from (D). Error bar SD (n=3 experiments).

**Figure S2. Mis12 protein can also be used as a kinetochore localizer in Opto-SAC.**

Endogenous Mis12 was tagged with PIF3-EGFP, and cells with or without PhyB621-Mph1ΔN and SynPCB1.0 were streaked onto two YEA plates. One plate was incubated at room temperature with continuous far-red light (Opto-SAC OFF) and the other was incubated with continuous red light (Opto-SAC ON) for 2 days.

**Figure S3. The timing of cell division before turning on the Opto-SAC is not associated with the failure of SAC silencing.**

Cyclin B (Cdc13-mNG) fluorescence intensities of each cell were measured at the start of the cell cycle (a nuclear division of the mother cell) before Opto-SAC ON (red shaded area). There was no clear relationship between the timing of Opto-SAC activation and competency to exit mitosis after Opto-SAC OFF.

**Table S1. The fission yeast strain used in this study**

