## Supplemental Information for "Development of optogenetic tools to manipulate cell cycle checkpoints"

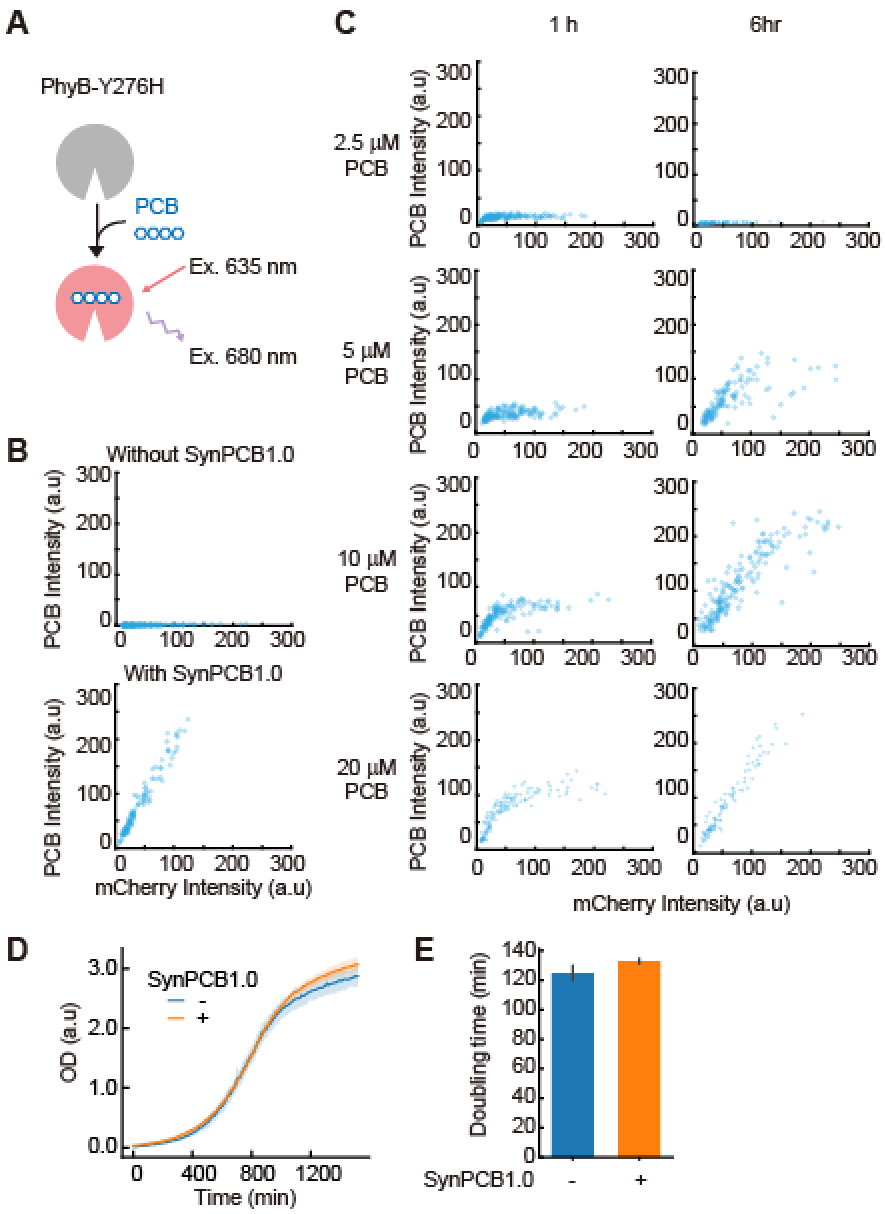


**Figure S1. Comparison between biosynthesis and uptake of PCB in *Schizosaccharomyces pombe*.** (A) Schematic of the measurement of PCB fluorescence. Infra-red PCB fluorescence is observed when PCB binds to the PhyB mutant PhyB-Y276H. (B) PCB fluorescence intensities in each cell are plotted as a function of mCherry fluorescence of PhyB-Y276H-mCherry in each cell. Upper and lower panels indicate control and SynPCB1.0-expressing cells, respectively. (C) PCB fluorescence intensities in each cell are plotted as a function of mCherry fluorescence of PhyB-Y276H-mCherry. Cells expressing PhyB-Y276H-mCherry were incubated with the indicated concentration of PCB for 1 h (left) and 6 h (right). At least 110 cells were quantified in each graph. (D) Growth curve of cells expressing SynPCB1.0. Solid lines indicate the mean OD from 3 independent cultures and shaded regions indicate the SD values. (E) Doubling time was calculated from (D). Error bar SD (n=3 experiments).

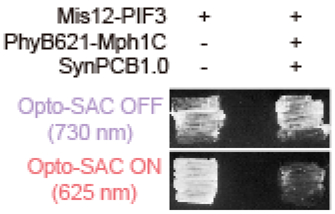


**Figure S2. Mis12 protein can also be used as a kinetochore localizer in Opto-SAC.**
Endogenous Mis12 was tagged with PIF3-EGFP, and cells with or without PhyB621-Mph1ΔN and SynPCB1.0 were streaked onto two YEA plates. One plate was incubated at room temperature with continuous far-red light (Opto-SAC OFF) and the other was incubated with continuous red light (Opto-SAC ON) for 2 days.

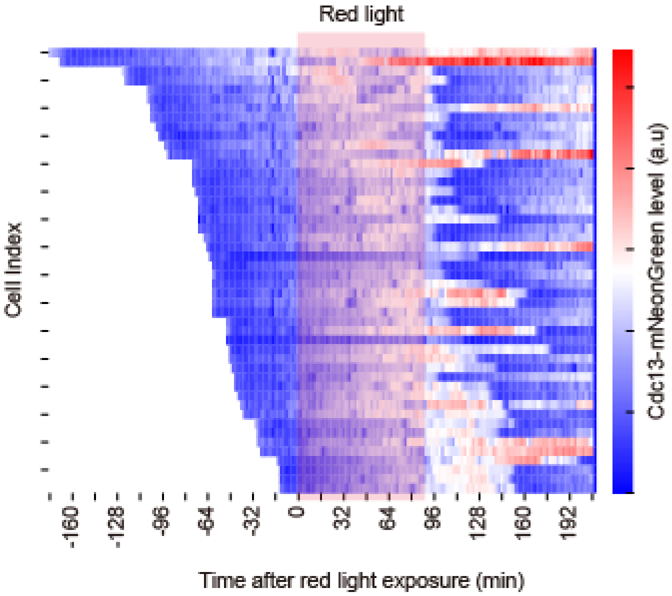


**Figure S3. The timing of cell division before turning on the Opto-SAC is not associated with the failure of SAC silencing.**

Cyclin B (Cdc13-mNG) fluorescence intensities of each cell were measured at the start of the cell cycle (a nuclear division of the mother cell) before Opto-SAC ON (red shaded area). There was no clear relationship between the timing of Opto-SAC activation and competency to exit mitosis after Opto-SAC OFF.

| Strain name | Genotype | origin | ref | Figure |
|---|---|---|---|---|
| YG074 | h+ ade6-M210 leu1-32 c1::Padh1-synPCB1.0<<kan | TN366 transformation | Uda et al., 2017 | - |
| YG153 | h+ ade6-M210 leu1-32 c1::Padh1-synPCB1.0<<kan z::Padh21-PIF3-EGFP<<nat | YG074 transformation | this study | - |
| YG151 | h+ ade6-M210 leu1-32 c1::Padh1-synPCB1.0<<kan z::Padh13-PIF3-EGFP<<nat | YG074 transformation | this study | - |
| YG200 | h+ ade6-M210 leu1-32 c1::Padh1-synPCB1.0<<kan c::Padh13-PIF3-EGFP<<hyg z::Padh1-hPhyB621-mCherry-ras1Δ200<<nat | YG153 transformation | this study | Fig 1 E,F |
| YG187 | h+ ade6-M210 leu1-32 c1::Padh1-synPCB1.0<<kan z::Padh13-PIF3-EGFP<<nat pREP1-PhyB621-mCherry-hta1 | YG151 transformation | this study | Fig 1 H,I |
| YG240 | h+ ade6-M210 leu1-32 c1::Padh1-synPCB1.0<<kan cdc2-PIF3-EGFP-hyg | YG074 transformation | this study | - |
| YG263 | h+ ade6-M210 leu1-32 c1::Padh1-synPCB1.0<<kan cdc2-PIF3-EGFP-hyg pREP1-PhyB621-mCherry-ras1deltaN200 | YG240 transformation | this study | Fig 2 B |
| YG333 | h+ ade6-M216 leu1-32 c1::Padh1-synPCB1.0<<bsd nuf2-PIF3-EGFP<<hyg z::Padh21-spPhyB621-mph1deltaC<<nat | YG332 transformation | this study | Fig 3 C,D,E, Fig 4, Fig S3 |
| YG332 | h+ ade6-M216 leu1-32 c1::Padh1-synPCB1.0<<bsd nuf2-PIF3-EGFP<hyg | YG076 transformation | this study | Fig 3 C |
| YG076 | h+ ade6-M216 leu1-32 c1::Padh1-synPCB1.0<<bsd | TN366 transformation | Uda et al., 2017 | |
| YG395 | h+ ade6-M216 leu1-32 c1::Padh1-synPCB1.0<<bsd nuf2-PIF3-EGFP<<hyg z::Padh21-spPhyB621-mph1deltaC<<nat mad2::kan | YG333 transformation | this study | Fig 3 C |
| YG90 | h- leu1? z1::Padh1-PhyB621-Y276H-mCherry<<bsd c1::Padh1-synPCB1.0<<kan | YG73xYG84 | Uda et al., 2017 | Fig S1 |
| YG331 | h+ ade6-M216 leu1-32 c1::Padh1-synPCB1.0<<bsd mis12-PIF3-EGFP<<hyg | YG076 transformation | this study | Fig S2 |
| YG335 | h+ ade6-M216 leu1-32 c1::Padh1-synPCB1.0<<bsd mis12-PIF3-EGFP<<hyg z::Padh21-spPhyB621-mph1deltaC<<nat | YG331 transformation | this study | Fig S2 |

**Table S1. The fission yeast strain used in this study**
